# The 2019-new coronavirus epidemic: evidence for virus evolution

**DOI:** 10.1101/2020.01.24.915157

**Authors:** Domenico Benvenuto, Marta Giovannetti, Alessandra Ciccozzi, Silvia Spoto, Silvia Angeletti, Massimo Ciccozzi

**Affiliations:** Unit of Medical Statistics and Molecular Epidemiology, University Campus Bio-Medico of Rome, Italy; Laboratório de Flavivírus, Instituto Oswaldo Cruz, Fundação Oswaldo Cruz, Rio de Janeiro, Brazil; Internal Medicine Unit, University Campus Bio-Medico of Rome, Italy; Unit of Clinical Laboratory Science, University Campus Bio-Medico of Rome, Italy

**Keywords:** 2019-nCoV, Molecular Epidemiology, Phylogeny, SARS

## Abstract

There is concern about a new coronavirus, the 2019-nCoV, as a global public health threat. In this article, we provide a preliminary evolutionary and molecular epidemiological analysis of this new virus. A phylogenetic tree has been built using the 15 available whole genome sequence of 2019-nCoV and 12 whole genome sequences highly similar sequences available in gene bank (5 from SARS, 2 from MERS and 5 from Bat SARS-like Coronavirus). FUBAR analysis shows that the Nucleocapsid and the Spike Glycoprotein has some sites under positive pressure while homology modelling helped to explain some molecular and structural differences between the viruses. The phylogenetic tree showed that 2019.nCoV significantly clustered with Bat SARS-like Coronavirus sequence isolated in 2015, whereas structural analysis revealed mutation in S and nucleocapsid proteins. From these results, 2019nCoV could be considered a coronavirus distinct from SARS virus, probably transmitted from bats or another host where mutations conferred upon it the ability to infect humans.

---

The family *Coronaviridae* comprises large, single, plus-stranded RNA viruses isolated from several species and previously known to cause common colds and diarrheal illnesses in humans [1]. A previously unknown severe acute respiratory syndrome (SARS) was observed in 2002 and later in 2003, a new coronavirus (SARS-CoV) was associated with the SARS outbreak [1]. Recently, a new coronavirus (2019-nCoV), has emerged in the region of Wuhan (China) as a cause of severe respiratory infection in humans. A total of 440 pneumonia cases have been confirmed in China up today (the State Council Information Office in Beijing, capital of China, Jan. 22, 2020). Animal to human transmission is considered the origin of epidemics as many patients declared to have visited a local fish and wild animal market in Wuhan in November. Quite recently, evidence has been gathered for animal to human and inter-human transmission of the virus [2][3].

While prompt diagnosis and patient isolation are the hallmarks for an initial control of this new epidemic, molecular epidemiology, evolutionary models and phylogenetic analysis can help estimate genetic variability and the evolutionary rate that in turn have important implications for disease progression as well as for drug and vaccine development. In this short report, we provide a phylogenetic three of the 2019 nCoV and identify sites of positive or negative selection pressure in distinct regions of the virus.

The complete genomes of 15 2019-nCoV sequences have been downloaded from GISAID (https://www.gisaid.org/) and GenBank (http://www.ncbi.nlm.nih.gov/genbank/). A dataset have been built using 5 Highly similar sequences for Severe Acute Respiratory Syndrome Virus (SARS), 2 sequences for Middle East Respiratory Syndrome (MERS) and 5 Highly similar sequences for Bat Severe Acute Respiratory Syndrome Virus respectively. The percentage of similarity has been identified using Basic Local Alignment Search Tool (BLAST; https://blast.ncbi.nlm.nih.gov/Blast.cgi), eventual duplicated sequences have been excluded from the datasets. The Dataset including 27 sequences has been aligned using multiple sequence alignment (MAFFT) online tool [4] and manually edited using Bioedit program v7.0.5 [5].

Maximum likelihood (ML) methods were employed for the analyses because they allow for testing different phylogenetic hypotheses by calculating the probability of a given model of evolution generating the observed data and by comparing the probabilities of nested models by the likelihood ratio test. The best-fitting nucleotide substitution model was chosen by JModeltest software [6]. ML tree was reconstructed using Generalised time reversible (GTR) plus Gamma distribution and invariant sites (+G +I) as evolutionary model using MEGA-X. [7]

Adaptive Evolution Server (http://www.datamonkey.org/) was used to find eventual sites of positive or negative selection. At this purpose the following tests has been used: fast-unconstrained Bayesian approximation (FUBAR) [8]. These tests allowed to infer the site-specific pervasive selection, the episodic diversifying selection across the region of interest and to identify episodic selection at individual sites [9]. Statistically significant positive or negative selection was based on p value < 0.05 [9].

Homology models have been built relying on the website SwissModel [10]. Structural templates have been searched and validated using the software available within the SwissModel environment and HHPred [11]. Homology models have been validated using the QMEAN tool [12]. Three-dimensional structures have been analyzed and displayed using PyMOL [13]. To map the structural variability of the N, E, S and M regions of the virus and their sites under selection pressure, homology modelling has been applied on the sequence of 2019-nCoV.

The ML phylogenetic tree, performed on whole genome sequences, is represented in Figure 1. In the tree, MERS virus sequences formed a distinct clade (clade I) from Bat SARS-like coronavirus, SARS virus and the 2019-nCoV clustering together in clade II. This clade includes two different clusters, the cluster IIa with Bat SARS-like coronavirus and the 2019-nCoV sequences whereas the cluster IIb with the Bat SARS-like coronavirus and the SARS virus sequences. The 2019-nCoV is significantly closely related only to the specific Bat SARS-like coronavirus isolated from Rhinolophus sinicus in 2015 in China (MG772934.1) (Figure 1).

**Figure 1.**
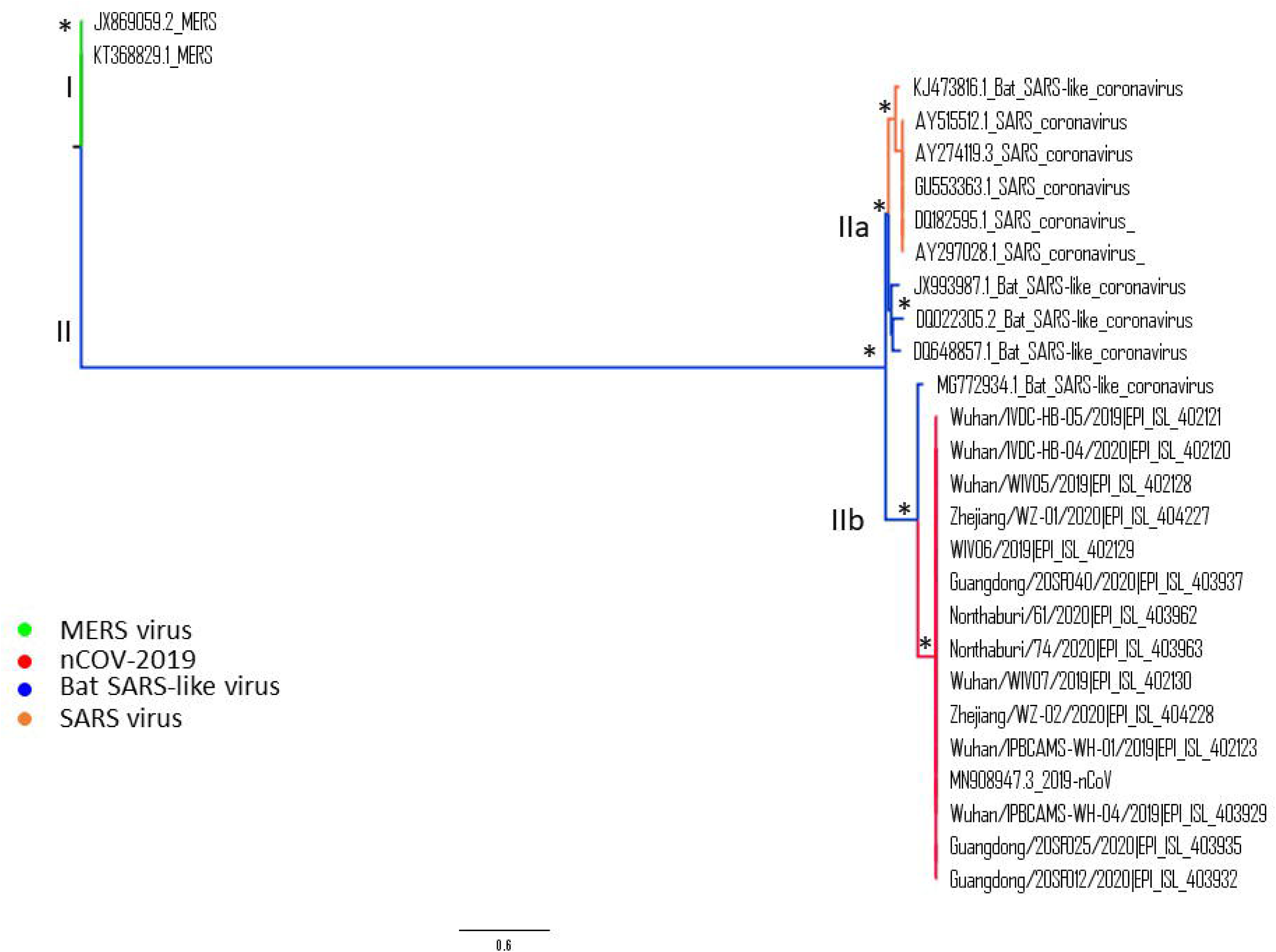
Maximum likelihood tree of 2019-nCoV. Along the branches an asterisk (*) indicating a statistical value from bootstrap (>90%).

Regarding the FUBAR analysis performed on the N Region, significant (p-value<0.05) pervasive episodic selection was found in 2 different sites (380th and 409th nucleotide position using the reference sequence; Wuhan seafood market pneumonia virus isolate labeled Wuhan-Hu-1 MN908947.3). Concerning the sequences in clade II, on the 409th aminoacidic position in Wuhan coronavirus sequence there is a glutamine instead of an asparagine residue while on the 380th aminoacidic position in Wuhan coronavirus sequence there is a threonine residue instead of an alanine.. Significant (p < 0.05) pervasive negative selection in 6 sites (14%) has been evidenced and confirmed by FUBAR analysis.

On the S Region, significant (p-value < 0.05) pervasive episodic selection was found in 2 different sites (536th and 644th nucleotide position using the reference sequence; Wuhan seafood market pneumonia virus isolate labeled Wuhan-Hu-1 MN908947.3). For the sequences in clade II, on the 536th aminoacidic position in Wuhan coronavirus sequence there is a asparagine residue instead of an aspartic acid residue while on the 644th aminoacidic position in Wuhan coronavirus sequence there is a threonine residue instead of an alanine residue. Significant (p < 0.05) pervasive negative selection in 1065 sites (87%) has been evidenced and confirmed by FUBAR analysis, suggesting that S Region could be highly conserved.

No sites under positive selection sites have been found in the E and M region.

The N region of the 2019-nCoV homology model has been built using a SARS coronavirus nucleocapsid protein structure (2jw8.1) since the statistical test have shown that this is the most stable and similar model among the other possible structures (Ramachandran Favoured 90.52%, GMQE 0.17, QMEAN -2.43), while for the S region the 2019-nCoV homology model has been built using a SARS coronavirus spike glycoprotein (6acc.1) for the same reasons (Ramachandran Favoured 87.87%, GMQE 0.83, QMEAN -3.14)

The further structural and molecular analysis of the Nucleocapsid region of the 2019-nCoV (MN908947.3) has highlighted that 2019-nCoV and the Bat SARS-like coronavirus (MG772934.) share the same aminoacidic sequence near the 309th position (SKQLQQ) while the SARS reference genome has a different aminoacidic sequence (SRQLQN). The same results have been found in the 380th aminoacidic position (KADET for 2019-nCoV and Bat SARS-like coronavirus and KTDEA for the SARS reference genome), in particular, in this case, the 2019-nCoV has a polar amino acid while the SARS has a non-polar amino acid (Figure 2).

**Figure 2.**
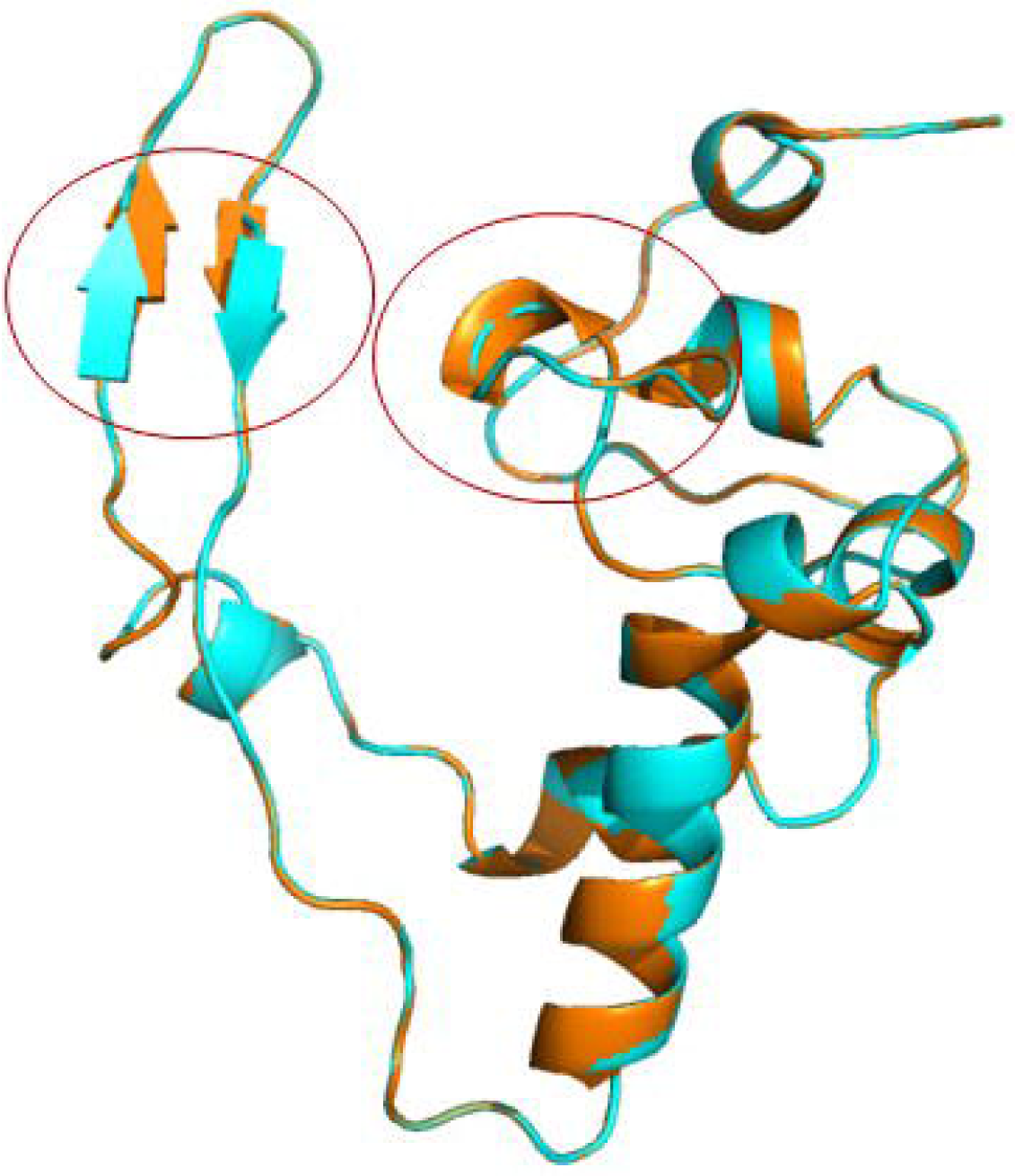
Cartoon model of the structural superposition between the homology model of the 2019-nCoV in blue and the Nucleocapsid protein of SARS Coronavirus (PDB code 2jw8.1) in orange. the presence of an alpha-helix on the SARS-CoV and not present on the 2019 n-CoV structure and e positional difference of the beta-sheets.

**Figure 3.**
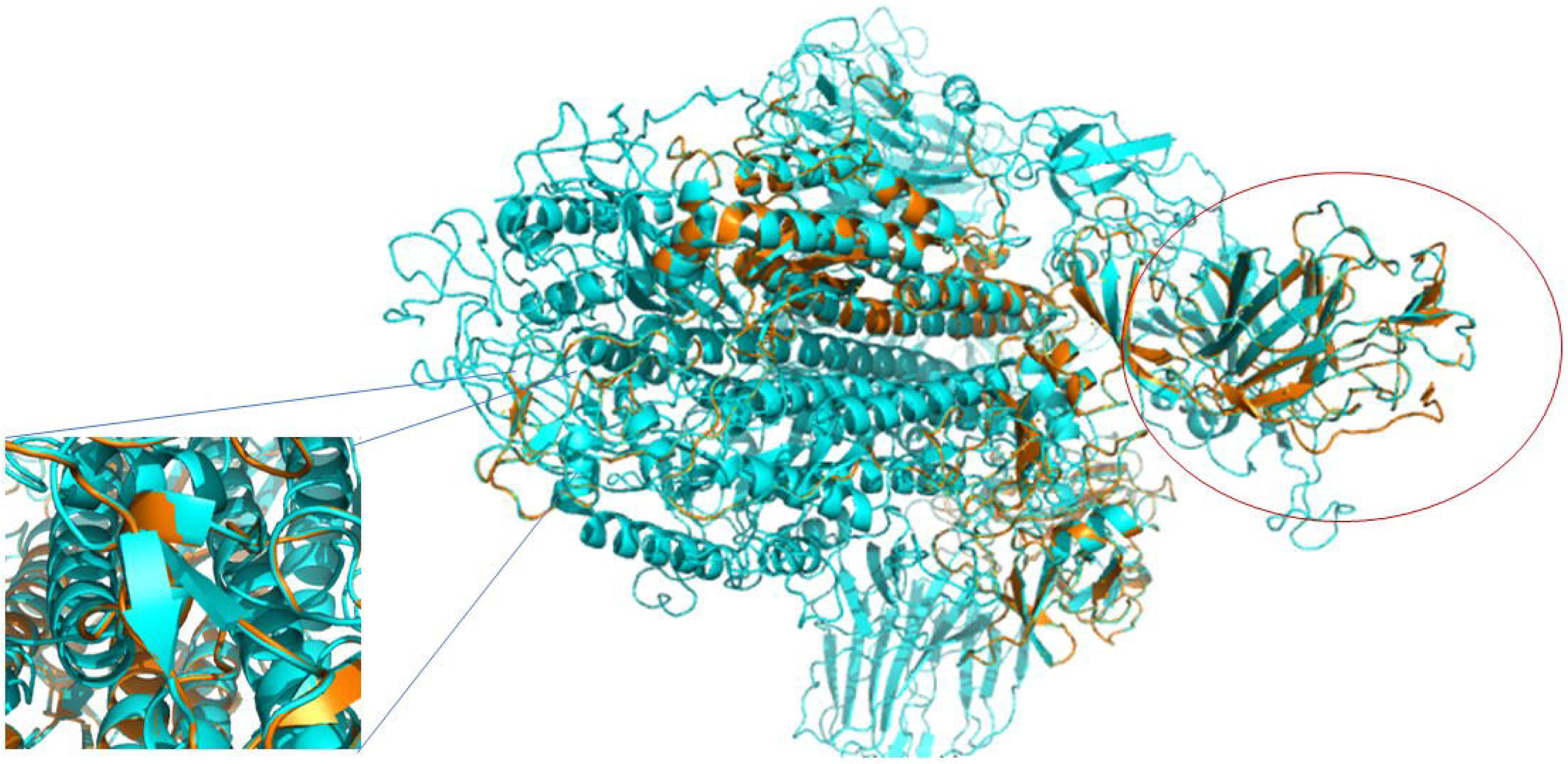
Cartoon model of the structural superposition between the homology model of the 2019-nCoV in blue and the spike glycoprotein of SARS Coronavirus (PDB code 6acc.1) in orange. the red circle highlights the presence of a variable region on the 2019 n-CoV at the beginning of the protein while the blue square highlights the presence of 2 beta-sheets on the 2019 n-CoV (401:KYR and 440:LND) that are not present on the SARS-CoV structure.

Regarding the sites under positive selective pressure found on the Spike Glycoprotein, the results have shown that the 536th aminoacidic position in 2019-nCoV has an asparagine residue while the Bat SARS-like coronavirus has a glutamine residue, the SARS virus, instead, has an aspartic acid residue. On the 644th aminoacidic position in 2019-nCoV sequence there is a threonine residue while the Bat SARS-like virus has a serine residue, instead the SARS virus has an alanine residue The data reported above show that the new 2019-nCoV significantly clustered with a sequence from the Bat SARS-like Coronavirus isolated in 2015. Moreover, in the phylogenetic tree, these two sequences are separated from the other Bat SARS-like coronavirus sequences, suggesting that this Bat SARS-like coronavirus is homologous and genetically more similar to the 2019-nCoV than to the other sequences of Bat SARS-like coronavirus. This supports the hypothesis that the transmission chain began from bat and reached the human. All other genomic sequences represented in the phylogenetic tree, also including SARS and MERS coronavirus, clustered separately thus excluding that the virus involved in the actual epidemic could belong to these sub-genuses. The structural analysis of two important viral protein, the nucleocapsid and the spike-like nucleoprotein (protein S), confirmed the significant similarity the new coronavirus with the bat-like SARS coronavirus and its difference from SARS coronavirus.

From the selective pressure and structural analysis, mutations of surface proteins, as the spike protein S, and of nucleocapsid N protein conferring stability to the viral particle have been shown. The viral spike protein is responsible for virus entry into the cell after by binding to a cell receptor and membrane fusion, two key roles in viral infection and pathogenesis. The N protein is a structural protein involved in virion assembly, playing a pivotal role in virus transcription and assembly efficiency. Mutation of these proteins could determine two important characteristics of the coronavirus isolated during the 2019nCoV epidemic, a higher stability than the bat-like SARS coronavirus and a lower pathogenicity than SARS coronavirus. These features can explain the 2019-nCoV zoonotic transmission and its initial lower severity than SARS epidemic. These results doesn’t exclude that further mutation due to positive selective pressure, lead by the epidemic evolution, could favor an ehancement of pathogenicity and transmission of this novel virus.

Recently, Ji et al. described homologous recombination within the spike glycoprotein of 2019nCoV favoring cross-species transmission and suggested snake as probable virus reservoir for human infection because its Resampling Similarity Codon Usage (RSCU) bias is more similar to *Bungarus multicinctus* snake compared to other animals and humans[3]. In a previous article has been proven that compositional properties, mutation pressure, natural selection, gene expression and dinucleotides affect the codon usage bias of *Bungarus* species [14].

These data along with data from our study, enforcing bat origin of infection, could clarify the transmission dynamic of the 2019-nCoV supporting infection control policy during the ongoing epidemic.

