## Supplementary figures and images for "The 2019-new coronavirus epidemic: evidence for virus evolution"

### Supplementary Figure

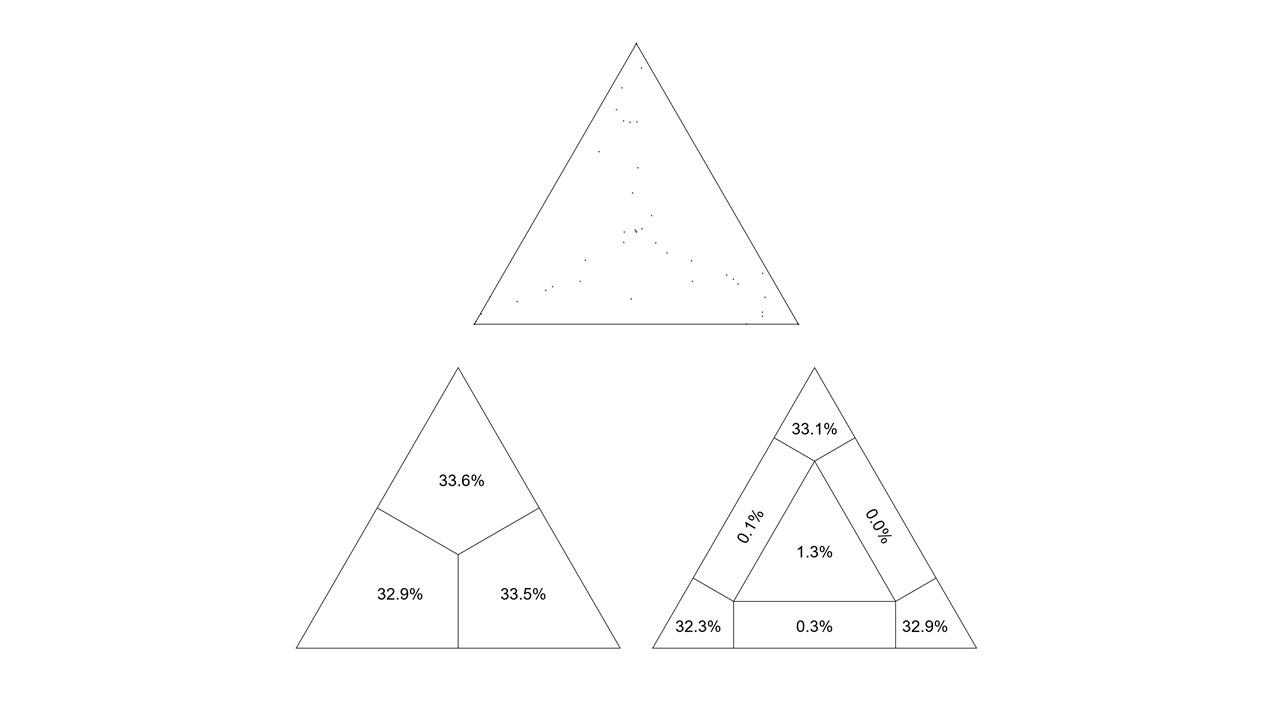
